# Applying Machine Learning to Increase Efficiency and Accuracy of Meta-Analytic Review

**DOI:** 10.1101/2020.10.06.314245

**Authors:** Aaron J. Gorelik, Mark G. Gorelik, Kathryn K. Ridout, Akua F. Nimarko, Virginia Peisch, Shamanth R. Kuramkote, Michelle Low, Tracy Pan, Simirthi Singh, Ananya Nrusimha, Manpreet K. Singh

## Abstract

The rapidly burgeoning quantity and complexity of publications makes curating and synthesizing information for meta-analyses ever more challenging. Meta-analyses require manual review of abstracts for study inclusion, which is time consuming, and variation among reviewer interpretation of inclusion/exclusion criteria for selecting a paper to be included in a review can impact a study’s outcome. To address these challenges in efficiency and accuracy, we propose and evaluate a machine learning approach to capture the definition of inclusion/exclusion criteria using a machine learning model to automate the selection process. We trained machine learning models on a manually reviewed dataset from a meta-analysis of resilience factors influencing psychopathology development. Then, the trained models were applied to an oncology dataset and evaluated for efficiency and accuracy against trained human reviewers. The results suggest that machine learning models can be used to automate the paper selection process and reduce the abstract review time while maintaining accuracy comparable to trained human reviewers. We propose a novel approach which uses model confidence to propose a subset of abstracts for manual review, thereby increasing the accuracy of the automated review while reducing the total number of abstracts requiring manual review. Furthermore, we delineate how leveraging these models more broadly may facilitate the sharing and synthesis of research expertise across disciplines.

## Background

A meta-analysis is a statistical methodology that quantitatively synthesizes data from individual studies, allowing overall trends to be determined for a research domain.^1^ Although meta-analyses are regarded as a strong systematic approach to review and synthesize evidence, they are often time- and labor-intensive, and limited by variations in researcher methodology, interpretation, and expertise.^2–4,11^ This is mainly due to the need for manual human review of studies for inclusion or exclusion criteria after an initial literature review using keyword searches from databases (e.g., Medline, PsyInfo, WebofScience). During this process of reviewing hundreds to thousands of abstracts, reviewers spend 1.5 minutes per abstract on average to decide which studies to initially include in the meta-analyses.^9,10^ The increasing rate of scientific publication across research domains expands the number of papers to be included in a given meta-analytic review, compounding this problem.^4,12,13^ Prior work has shown that applying machine learning (ML) approaches can select abstracts for meta-analytic inclusion or exclusion with increased efficiency and similar accuracy to human reviewers.^5–8^ However, these approaches have never been applied to new meta-analysis topics differing from the original training topic to see if the models can translate the same inclusion and exclusion criteria across research domains.^5–8^

Thus, we describe a ML-based protocol to improve efficiency relative to trained human reviewers by automating review of manuscript abstracts for inclusion/exclusion criteria. This protocol also leverages ML models to assist human reviewers in de novo manuscript selection for inclusion/exclusion, with the ability to modify the model sensitivity to achieve desired trade-offs between accuracy and time invested in manually reviewing abstracts. We conclude with a suggestion for the creation of meta-concept ML model repository to facilitate collaboration across research domains.

## Main

To accomplish the first goal of efficient automated review of abstracts for inclusion and exclusion criteria, the ML-models were first “trained and tested” and then “evaluated” for performance. The ML models tested were curated keywords search (search), Multinomial Naïve Bayes classifier, BERT (Bidirectional Encoder Representations from Transformers), and SciBERT (BERT trained on scientific literature). BERT and SciBERT models come pretrained using a self-supervised approach from a large text corpus;^14–16^ these were fine-tuned for each meta-concept (the summary of training and fine-tuning parameters and results for each ML model is summarized in Supplemental Table 1 and 2, respectively).

The “train and test” dataset contained 8202 abstracts selected for a meta-analysis examining resilience factors influencing psychopathology development in the field of psychiatry (PROSPERO protocol CRD42020172975 for details). The abstracts were evaluated for inclusion in or exclusion from four concept areas: Resilience, Biomarkers & Disease, Stressors, and Conditions. These concepts were defined using keywords and classification guidelines identified by experts in collaboration with librarians trained in systematic reviews (see Table 1 for details). Standard meta-analytic methods were used to train human reviewers to promote inter-rater agreement when reviewing for inclusion and exclusion for each concept area (see methods for training protocol).^3^ These standard training methods ensured accuracy of human review and confirmed labeling needed for training four different ML models.

**Table 1.** Meta-concepts

| Meta-Concept | Classification Guidelines | # Key Words | Example Common Terms |
| --- | --- | --- | --- |
| Resilience | The ability to adapt in the face of adversity and stress | 169 | Change, adapt, resilience, risk factors, adjustment |
| Biomarkers & Disease | Physical disorders or biological markers | 282 | Gene expression, oxidative stress, molecular, brain, RNA expression |
| Conditions | Psychiatric or neurological disorders; conditions related to brain health | 563 | Depression, psychopathology, anxiety, schizophrenia, internalizing |
| Stressors | Events or conditions in early life environment that may trigger stress | 253 | Stress, injury, death, trauma, SES |

**Table 2.**
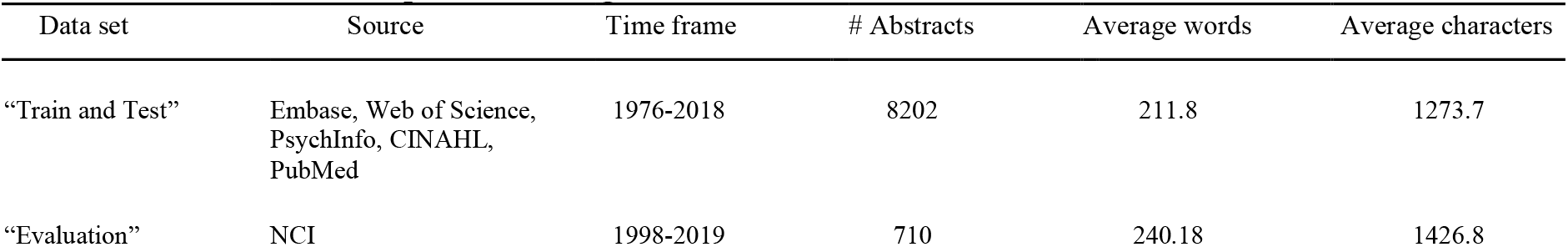
Datasets used for the process. Average words and characters refers to the words and characters in the abstracts.

| Data set | Source | Time frame | # Abstracts | Average words | Average characters |
| --- | --- | --- | --- | --- | --- |
| “Train and Test” | Embase, Web of Science, PsychInfo, CINAHL, PubMed | 1976-2018 | 8202 | 211.8 | 1273.7 |
| “Evaluation” | NCI | 1998-2019 | 710 | 240.18 | 1426.8 |

The “evaluation” dataset contained abstracts from the field of oncology to evaluate how accurately these ML concepts transferred to a new research domain (see Table 2 for details). Tagging by a domain and meta-analysis expert (KR) was used as the “ground truth.” Accuracy of the four different automation methods were compared against trained human reviewers and an untrained domain expert. The “evaluation” dataset was processed by automation, human reviewers, and the untrained domain expert without additional training. A subset of the abstracts in the “evaluation” dataset (*n*=360, 99% confidence with 5% margin of error) was compared against ground truth to ascertain relative performance. The overall process for using ML to automate abstract review is described in Figure 1.

**Figure 1:**
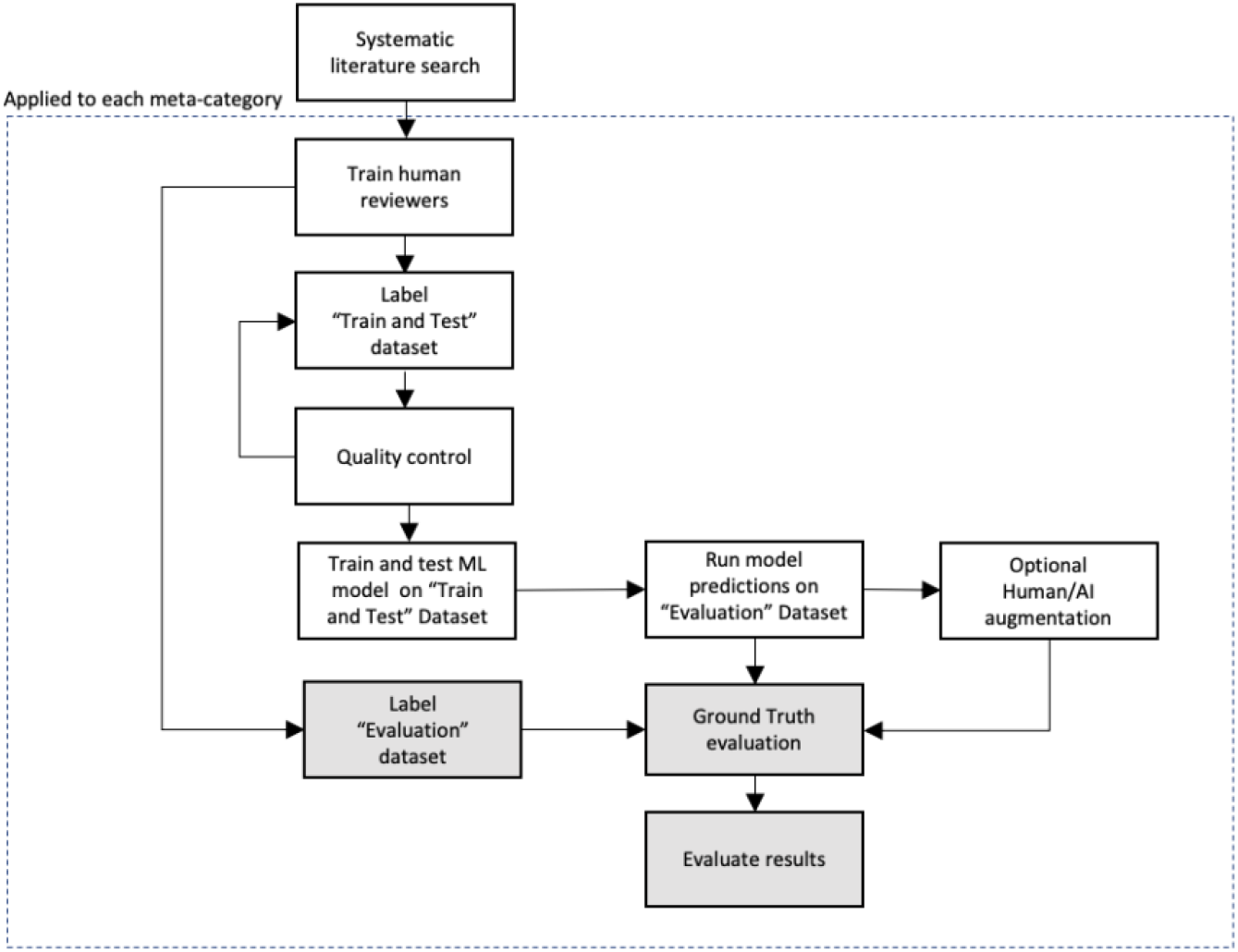
Process flow for individual meta-concept. “train and test” dataset was from the meta-analysis examining resilience factors influencing psychopathology development and the “evaluation” dataset was from NCI oncology dataset. The blocks in grey are only relevant to the evaluation of this methodology and would not be standard practice in real world application.

Of the four automation methods, SciBERT performed most accurately compared to human reviewers in the “evaluation” dataset, being marginally more accurate then trained human reviewers (+1% to +11%) and less accurate then the untrained expert (−3% to −11%). SciBERT had greater sensitivity for categorizing abstracts to the Biomarkers & Disease (+12.2%) and Resilience (+1.6%) meta-concepts and lower sensitivity for the Stressors (−14.3%) and Conditions (−22.5%) meta-concepts compared to the trained human reviewers (see Figure 2 for details). In addition, compared to human reviewers, SciBERT’s classification performance had better precision and recall categorizing abstracts to the Conditions (F_1_=+0.072), Biomarkers (F_1_=+0.096), and Resilience (F_1_=+0.12) meta-concepts, and slightly worse performance for the Stressors (−0.03) meta-concept (see Figure 2 for details).

**Figure 2:**
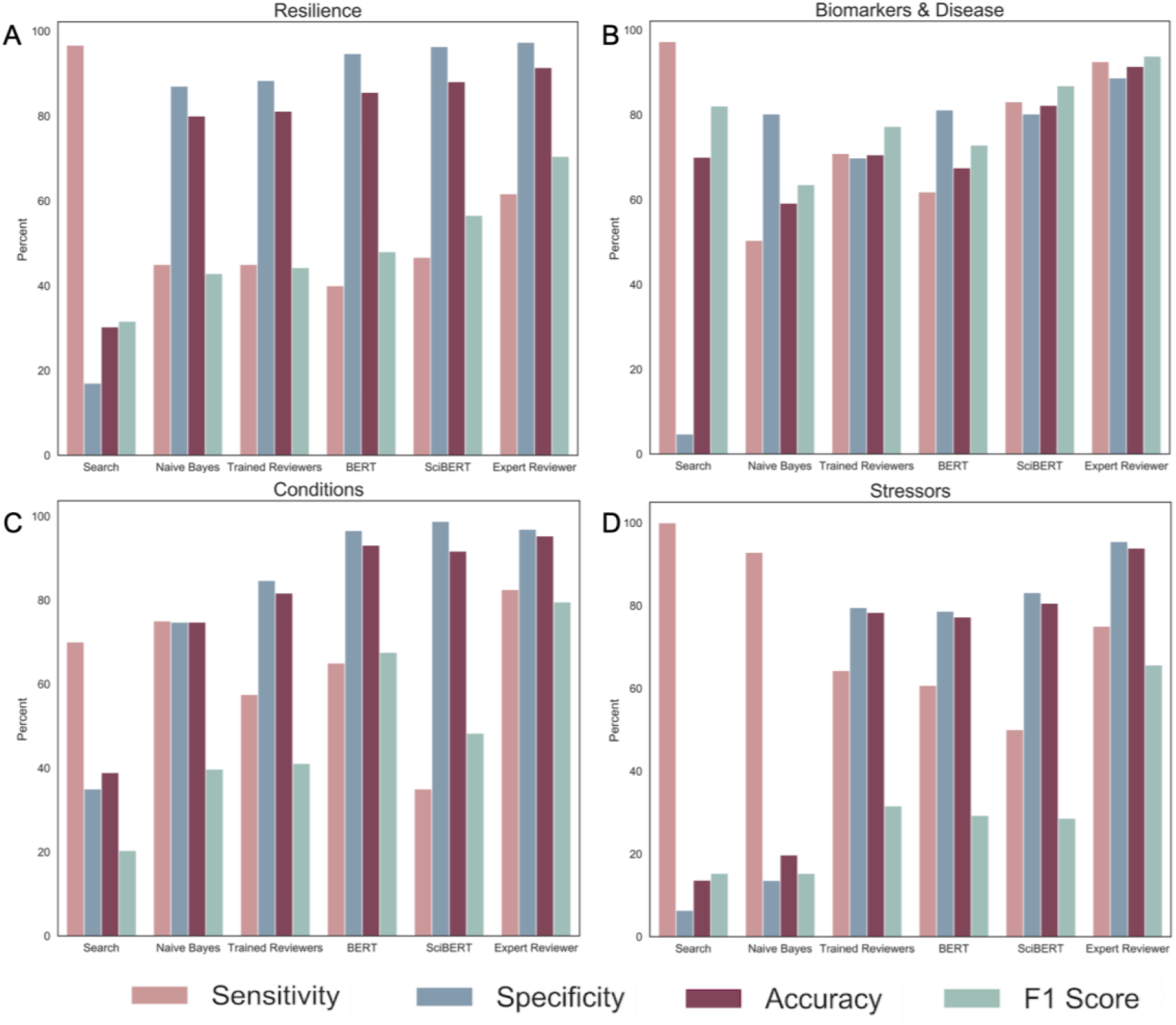
Paper Selection Methods Performance. Performance of all selection methods against ground truth for each meta-concept: A Resilience, B Biomarkers & Disease, C Conditions, D Stressors.

After finetuning, the SciBERT models took 0.075 seconds on average to infer inclusion/exclusion per abstract (see Table S4 for training loop and prediction times and methods for hardware details). This is compared to the trained human reviewers, who took 90 seconds on average to manually review an abstract. These results suggest that ML models trained on abstracts for specific inclusion and exclusion criteria can be translated to a new research domain while significantly reducing the time and effort required to review abstracts (Figure 2).

To accomplish the second goal of using ML models to assist human reviewers in efficient and accurate de novo abstract selection for inclusion/exclusion, we used SciBERT, which was the most similar to ground truth. SciBERT confidence scores for abstract non-inclusion *(1.0 - p(include)*) were used to recommend abstracts that needed additional human review (see Figure S3). Lower and higher SciBERT confidence ranges were tested for each concept area, and then examined for abstract classification accuracy versus number of abstracts requiring human review (please see methods for overview of process). For the low confidence range, abstracts flagged for manual review ranged from 3% of abstracts for the Conditions concept to 49% for the Stressors concept. When adding in the manual review, the accuracy and sensitivity increased by 7% and 12.5% respectively for the Conditions meta-concept, while for the Stressors meta-concept accuracy and sensitivity increased by 12% and 43%, respectively. For the high confidence range, abstracts flagged for manual review ranged from 6% for the Conditions concept to 73% for the Stressors concept. Combining the SciBERT and manual review, accuracy and sensitivity increased by 7% and 27.5% respectively for the Conditions concept, while accuracy and sensitivity increased by 15% and 46% for the Stressors concept, respectively. (see Figure 3 for all the results for both low human review and high sensitivity for each meta-concept for both augmentation types). As such, this ML approach can flag a subset of abstracts based on model confidence for manual review for inclusion/exclusion in meta-analyses, allowing human reviewers to calibrate between abstract review accuracy versus time spent manually reviewing abstracts.

**Figure 3:**
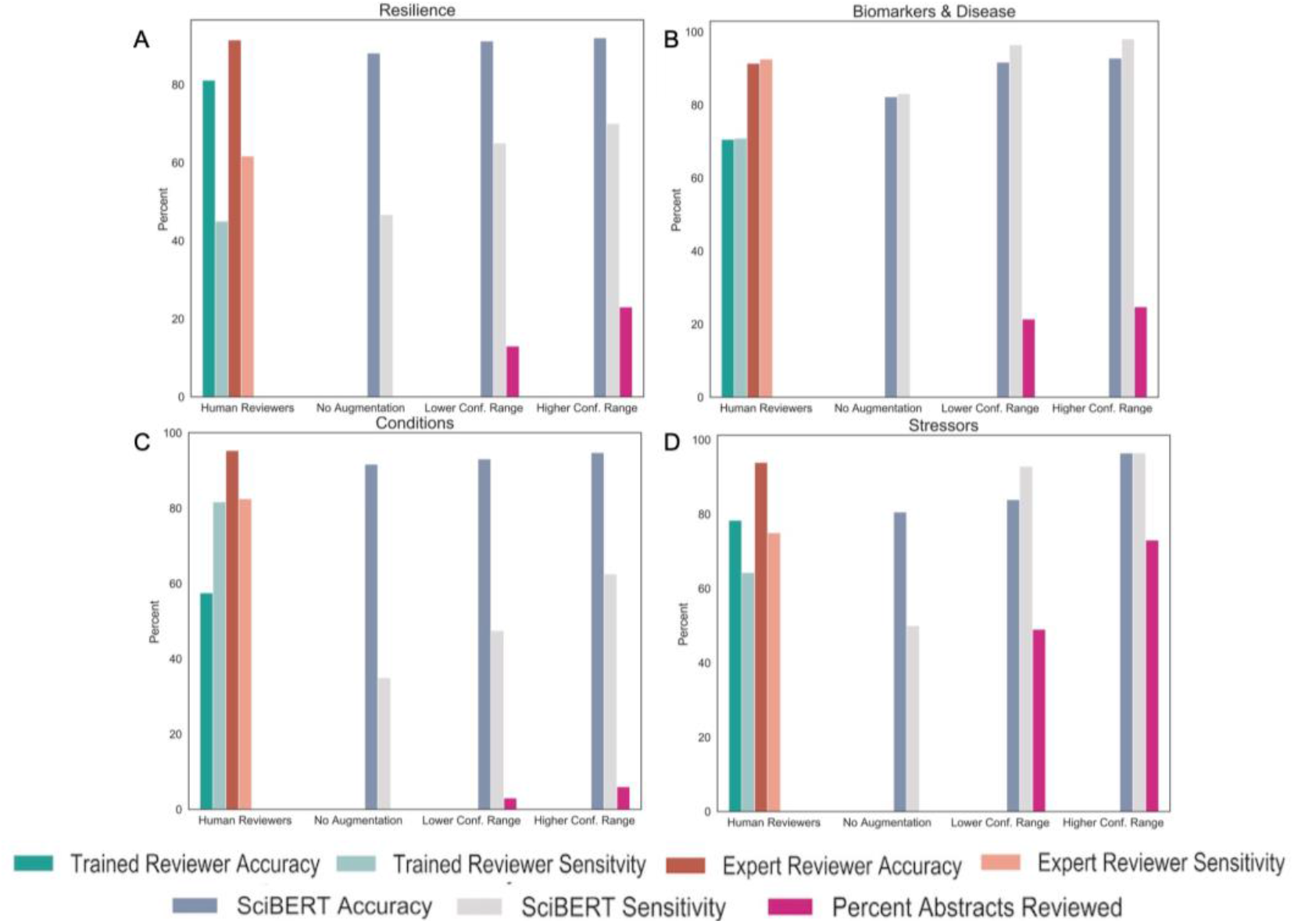
Human/AI augmentation performance. Shows use of artificial intelligence augmentation on volume of abstracts to review and impact on sensitivity and accuracy compared to trained human reviewers and untrained expert relative to ground truth for each meta-concept: A Resilience, B Biomarkers & Disease, C Conditions, D Stressors.

**Figure 4:**
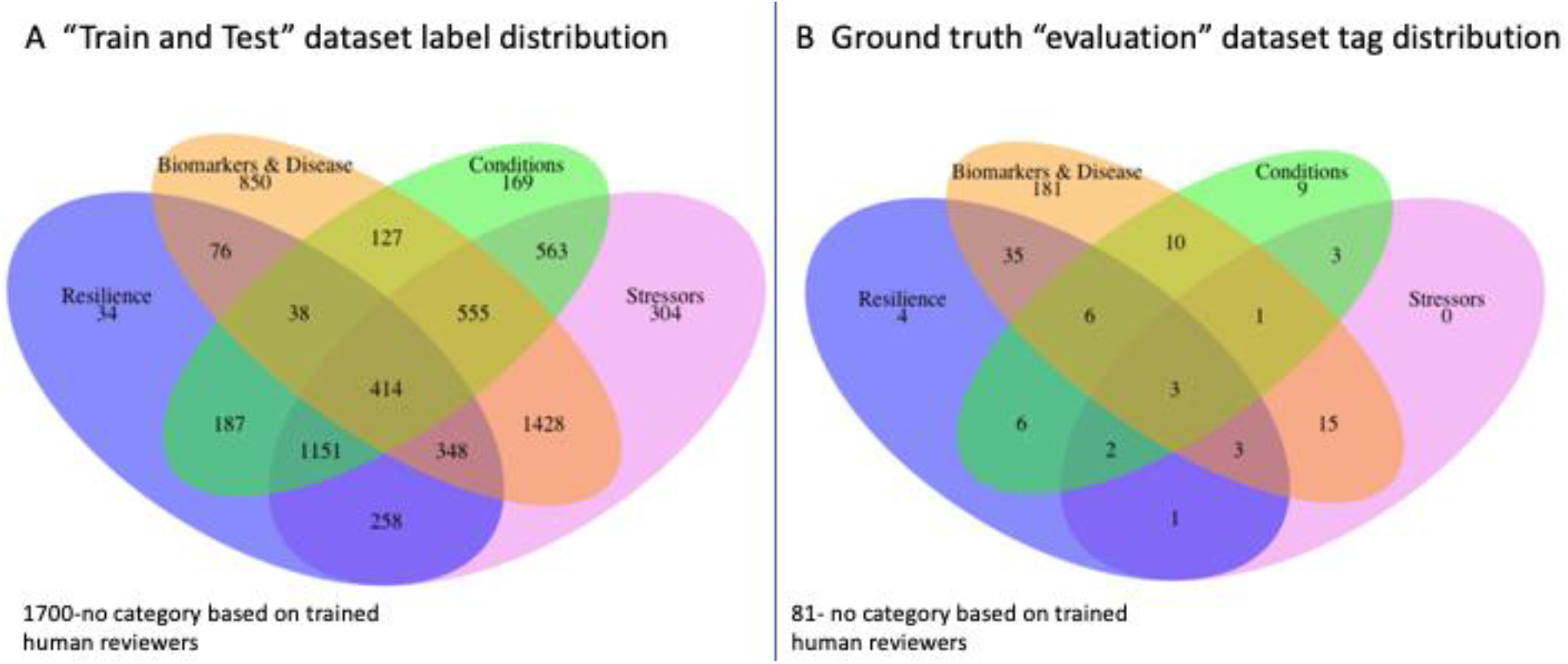
Meta-concept Overlaps. A. “train and test” dataset label distribution shows how *n*=8202 abstracts were manually tagged by trained human reviewers. B. “evaluation” dataset *n*=360 label distribution tagged by ground truth.

As the body of scientific knowledge grows and scientists become more specialized, evidence synthesis across disciplines is becoming increasingly challenging. Technology can be leveraged to help us organize and connect knowledge across domains. Prior work focused on similarity matching, which looks at clustering abstracts based on the similarity of words/sentences.^7,8^ Similarity matching works well for systematic reviews that have narrow inclusion criteria (e.g. specific terminology) but does not support the automated transfer of meta-concept to different domains. Consequently, prior models may be less useful for either reproducibility or generalizability for future meta-analytic reviews.^7,17,18^ BERT and SciBERT ML models leverage bidirectional representations to generate pretrained models drawing from a broader set of concepts. With refinement of the semantic and structural knowledge captured during the pretraining stages (transfer learning) to a substantially smaller data set, we were able to transfer ML model concepts across research domains (e.g. from psychiatry to oncology datasets).^14,15^ In addition, by creating a separate ML model for each concept, we are able to use the concepts as filters that can be utilized to find papers that have any combination of these concepts (See Figure 4). This feature is particularly useful for the repurposing of these models to future meta-analyses and sets the foundation for conceptualizing ML meta-concept repositories (See Figure 4 for meta-concept interactions in the evaluation *n*=360 and the “train and test” datasets).

We acknowledge the limitations of our current work. For our models, ground truth was assumed to always be accurate. However, human interpretation of concepts can introduce variation into a model. With the “evaluation” dataset, we did not retrain human reviewers regarding interpretation of the four concept areas to the new research domain, which is part of standard training to ensure inter-rater reliability (this is reflected in the trained reviewer error analysis, see supplement). However, not retraining human reviewers was done intentionally to evaluate the differences in human versus ML model interpretation of the concepts across the research domains.^19–21^ Further, BERT and SciBERT are computationally expensive and required truncation of text to 512 characters which could lead to potential data loss. This limitation is temporary as the most recent advancement in self-attention allows linear scaling with text size enabling our approach to apply to any size text.^22^ Finally, we acknowledge that model performance can vary for different datasets. Despite these limitations, we have created an ML-based method to augment meta-analysis review and have demonstrated practical improvements in accuracy and efficiency. Further, we have shown the possibility of transferring concepts across research domains to connect and expand knowledge across scientific fields.

## Future Direction

Our study provides an initial demonstration that advanced ML models can efficiently and accurately capture a meta-concept and apply the concept from a particular research field (e.g. psychiatry) to another research field (e.g. oncology). However, with these models, there is still significant work required to train reviewers, label data, and train or fine-tune a concept model. Creating a concept model repository with meta-concepts would facilitate sharing of these models and significantly reduce the efforts of researchers in creating, refining, sharing, and applying ML models across research fields, in order to easily and efficiently select papers that are relevant to meta-concepts. Additionally, this would also facilitate the creation of a common ontology that could be shared across different meta-analyses and across disciplines. For this process to be effective, standardization for creating and documenting meta-concept ML models would need to be developed. Some of the key components for this process, such as defining and evaluating meta-concept models have been described in the methods section.

Our approach demonstrates the feasibility of applying ML to meta-analytic methods to improve efficiency and accuracy within and across research fields. A repository of ML models to represent different meta-concepts could enable researchers to more broadly share and synthesize their expertise to integrate knowledge across scientific fields.

## Author Contributions

AG, MG, KR, and MS conceptualized and executed the study protocol. AG, MG, and KR contributed to the analyses. AG, MG, MS, KR, VP, and AFN contributed to writing of the manuscript. AG and MG contributed to visualizations. KR, AN AFN, VP, SK, TP, ML, SS, and MS contributed to the manual tagging of abstracts.

## Code availability

The code, models, and datasets are available at https://github.com/MetaAnalysisPipeline

## Acknowledgments

We would like to express appreciation to the members of our Pediatric Emotion And Resilience Lab (PEARL) for their contributions to this work.

## Methods

### Meta-Analysis Protocol and Registration

The meta-analysis examining resilience factors influencing psychopathology development protocol was registered with the International Prospective Register of Systemic Reviews (PROSPERO, CRD42020172975). The study criteria were designed using the Preferred Reporting Items for Systematic Reviews and Meta-Analyses (PRISMA). Portions of The Meta-analysis Of Observational Studies in Epidemiology (MOOSE) Guidelines were also followed and adapted into PRISMA.^23,24^

### “Training” meta-analysis study eligibility

Studies were included for the meta-analysis examining resilience factors influencing psychopathology development if they (1) examined the effects of early adversity in the form of abuse, neglect, socioeconomic status (SES), or other adverse exposures on human subjects occurring prenatally up to age 18; and (2) provided adequate description of adversity assessments. Studies using indirect proxies of early adversity, such as parental education alone, were excluded. Prospective, observational, and retrospective studies were considered.

### Meta-Analysis Information Sources and Search Strategy

For the meta-analysis examining resilience factors influencing psychopathology development, a comprehensive electronic search conducted in September 2018 identified English language studies indexed in PubMed/Medline, PsycINFO, CINAHL, and Web of Science; no publication date limitation was set. The search was performed by investigators with topic clinical and research experience (KR) in consultation with a librarian trained in systematic reviews. The search strategy included terms and combinations to identify early life stress, biomarkers and resilience (See Supplementary Table 3 for search criteria). Primary study and review article references were searched; studies were appraised for inclusion or exclusion using an a priori criteria as described under study eligibility.

### Datasets

Two different abstract datasets were utilized: a “train and test” data set for training the ML models (obtained from the meta-analysis examining resilience factors influencing psychopathology development) and an “evaluation” data set (oncology research field) to evaluate the ML models’ performance (see Table 2 for dataset details). For the “evaluation” (oncology) dataset, we used the NCI cancer control publications from 1995-2019 where author type included was “Both NCI & Extramural Researchers.”

### Concept inclusion and exclusion criteria

A definition was created for each of the four concepts based on key terms developed for the “train and test” meta-analysis (see Table 1) and expanded by content experts (MKS and KR). For the “training” dataset, human reviewers were trained regarding inclusion and exclusion criteria for each concept as found in the reviewer training and dataset labeling section below. For the “evaluation” dataset, ground truth, the human reviewers, and the untrained domain expert were asked use their “judgment and intuition” regarding the four concepts in addition to searching the abstracts for the presence of key terms. Table 1 describes the criteria for each meta-concept (see supplement table 4a-4d for list of key words).

### Reviewer Training and dataset labeling

The manual review team (SK, ML, AFN, AN, TP, VP, and SS) consisted of members from varying degrees of domain expertise, including undergraduate, graduate, postdoctoral, and faculty level coders.^25^ After a group training regarding the goals of the study, the study inclusion/exclusion criteria, and concepts, the team tagged 150 identical abstracts obtained from the “train and test” data set, and then compared their results to ground truth. Any discrepancies in labeling were resolved in a joint training session. This training resulted in >90% inter-rater reliability. In addition, the reviewers went through an extensive training protocol including a quality control (QC) methodology to ensure consistent results (see QC section). Reviewers labeled both the “train and test” and “evaluation” abstract datasets. A research field expert and expert in meta-analysis methodology (KR) was used as the ground truth in validating the manual tagging results from the “train and test” dataset for abstracts where there was a difference in labeling between trained reviewer and ground truth, and as a final arbiter for any ambiguity. MKS, our untrained domain expert reviewer, did not undergo the aforementioned reviewer training or QC process.

The QC process and reviewer training was applied only to the “train and test” data sets. This was to allow for evaluation and comparison of meta-concept translation by humans compared to the ML models.

### Quality Control (QC)

The unlabeled abstracts in the “train and test” dataset were divided equally among the trained reviewers. Once a human reviewer had labeled their portion (~1,350 abstracts), 150 random abstracts were checked against ground truth (99% confidence with 5% margin of error). If the percent agreement rate with ground truth was below 90%, that reviewer’s portion was relabeled by two new reviewers after which 150 samples (99% confidence with 5% margin of error) were re-checked against ground truth. Only one reviewer’s result was below 90% agreement and was therefore relabeled and re-validated. In addition to QC, any reviewer could discuss abstracts in the “train and test” data set to the ground truth to resolve ambiguity. However, in the “evaluation” dataset (not part of the standard meta-analysis protocol), these QC protocols were not utilized to allow comparison of meta-concept application between human reviewers and the ML model.

### Automated Filters

Four different automated approaches for selecting papers were tested and validated: one keyword search and three machine learning based models (Multinomial Naive Bayes, BERT, SciBERT).^14,15,26^

Multinomial Naïve Bayes is a commonly used text classification approach that has been shown to perform well in applications such as spam filtering.^27^ We used the scikitlearn implementation of Naïve Bayes due to its widespread use. The Naïve Bayes model was configured to get best performance for each meta-concept. The model was trained on a term frequency (TF) normalized matrix of word counts from the training set for Stressors and Conditions meta-concepts. For Resilience and Biomarkers & Disease meta-concepts the model was trained on term frequency–inverse document frequency (TF-IDF) normalized matrix of word counts from the training set (for details of other parameters see Table S1 in supplement). The model was optimized using a grid search.

BERT and SciBERT have proven to be extremely powerful for picking up context dependent patterns that may be missed by more traditional approaches, such as Naïve Bayes. BERT and SciBERT leverage bidirectional representations to generate pretrained models. BERT and SciBERT both came pre-trained on vocabularies of approximately 30,000 unique words and subwords. The main difference between the two is that BERT is trained on general purpose text corpus (book corpus and Wikipedia), whereas SciBERT is trained on scientific literature.^14^ The texts used to pre-train BERT and SciBERT share approximately 42% of the unique words. All four meta-concepts used the same parameters (see table S2 in supplement). Pre-trained BERT and SciBERT models were downloaded from huggingface.^28^

All ML based models were trained and measured on the same training test split of the “train and test” dataset implemented in sklearn using a seed value of 42 to ensure reproducible results. The split was 90% training and 10% test data. A smaller test portion was chosen since the real validation was done against a separate “evaluation” dataset. The Multinomial Naïve Based model was trained from scratch, whereas BERT and SciBERT were fine-tuned on top of pretrained models based on different text corpuses. For fine-tuning, the abstract texts were truncated at 512 words (see supplement Figure 1 and 2 for details).

For keyword search testing, the keyword lists were constructed by domain experts for the keyword search testing model. All keywords were at least 3 characters and abstracts were included if they contained at least one keyword. Special characters, such as asterisks, were removed from the search terms in order to mitigate inconsistent usage.

BERT and SciBERT were fine-tuned and used on the Google Colab Pro environment using a Tesla P100-PCIE-16GB. All other code was executed on a MacBook Pro retina with 2.9 GHz i9 CPU.

### Combining Multiple Meta-Concepts

By creating a separate model for each concept, we were able to evaluate how often these concepts were studied together or individually. In effect, the concept became a search criterion that could be combined in a logical expression for searching for abstracts that span different categories (e.g. includes concept Resilience but not Biomarkers & Disease or Conditions). Thus, this enables researchers to use one dataset to confirm multiple possible combinations of concepts.

### Result Analysis

All abstracts in the evaluation data set were labeled using automated filtering methods. Accuracy, recall, and F_1_ score for all automated filtering approaches were calculated using sklearn implementation. Vectors were extracted from both the fine-tuned BERT and SciBERT models and dimensionality reduction to demonstrate generalizability using transfer learning approach was performed by UMAP (see supplement figure 4).^29^

All abstracts in the evaluation data set were labeled by a group of trained human reviewers. In addition, 360 random abstracts were selected from the “evaluation” dataset to achieve 99% confidence with 5% margin of error accuracy. The abstracts were labeled by an untrained expert and ground truth.

The results for 360 random abstracts were compared across all the different filtering/labeling methods for each meta-concept topic against ground-truth: untrained expert, trained human reviewers, key word search, and ML models: Naive Bayes, BERT and SciBERT. As an extra cautionary step, discrepancy in the 360 labels between ground truth and untrained domain expert were reviewed to reduce potential for ground truth errors. We used the following metrics to evaluate performance of each method:

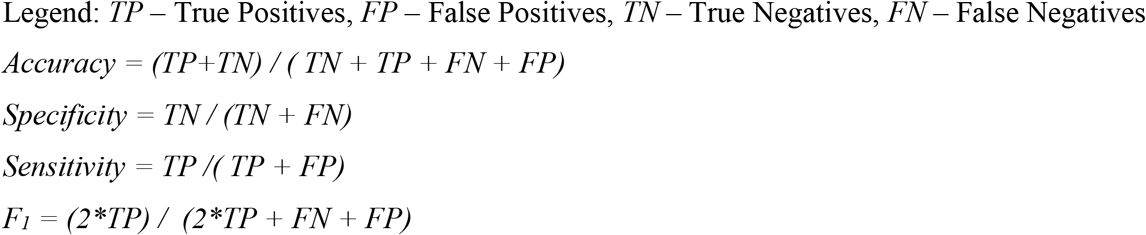

### AI/Human Augmentation

To enable the researchers to achieve a higher level of accuracy and sensitivity than a pure automated abstract selection process, the system generated a recommended list of abstracts to review. We tested low and high confidence ranges (defined below). This methodology was applied to the SciBERT model only, although the same methodology could be extended to BERT and Naïve Bayesian Classifier.

The ML model chose abstracts for manual human review within a range of negative class inference confidence values (*1-p(inclusion)*). For the lower confidence range, we used confidence values between 0.5 and 0.9, and for the higher confidence range we used values between 0.5 and 0.95. The abstracts with confidence below 0.5 were treated as true positives and values above 0.9 or 0.95 were treated as true negatives. For deep neural networks, the confidence values are typically clustered at the higher end of the confidence score (*p(inclusion)*)^30^ We chose to focus on sensitivity as the key evaluation in addition to accuracy. This also aligned well with the meta-analysis process where false positives would likely be weeded out at later stages of a meta-analysis.

We used ground truth for the evaluation dataset with the assumption that a human reviewer would always label the abstracts accurately for the augmentation step. We chose not to incorporate the accuracy of ground truth because the error for human reviewers depends on quality control applied and in a meta-analysis, which would be considered ground truth.

## Supplementary Methods & Extended Figures

### Multinomial Naïve Bayes

For any configuration values for scikitlearn package not listed in Table S1, the default values that came with the package were used.^1^

**Table S1.**
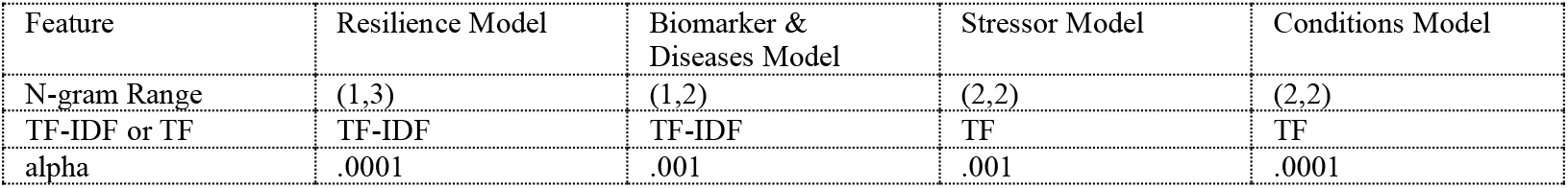
Final Naïve Bayes Model Parameters

| Feature | Resilience Model | Biomarker & Diseases Model | Stressor Model | Conditions Model |
| --- | --- | --- | --- | --- |
| N-gram Range | (1,3) | (1,2) | (2,2) | (2,2) |
| TF-IDF or TF | TF-IDF | TF-IDF | TF | TF |
| alpha | .0001 | .001 | .001 | .0001 |

### BERT & SciBERT

The parameters used to fine-tune the BERT and SciBERT models were the default recommendations based on the authors of BERT (Table S2).^2,3^ For any parameters not listed in Table S2, the default values were used that came with the huggingface package.^4^

**Table S2.** Final BERT and SciBERT model parameters

| Parameters | BERT | SciBERT |
| --- | --- | --- |
| Model | bert-base-uncased | allenai/scibert_scivocab_uncased |
| Learning Rate | $2e^{-5}$ | $2e^{-5}$ |
| Adam Epsilon | $1e^{-4}$ | $1e^{-4}$ |
| Batch size | 8 | 4 |
| Epochs | 4 | 4 |
| Text Truncation Cutoff | 512 | 512 |
| Seed | 42 | 42 |

**Fig S1.**
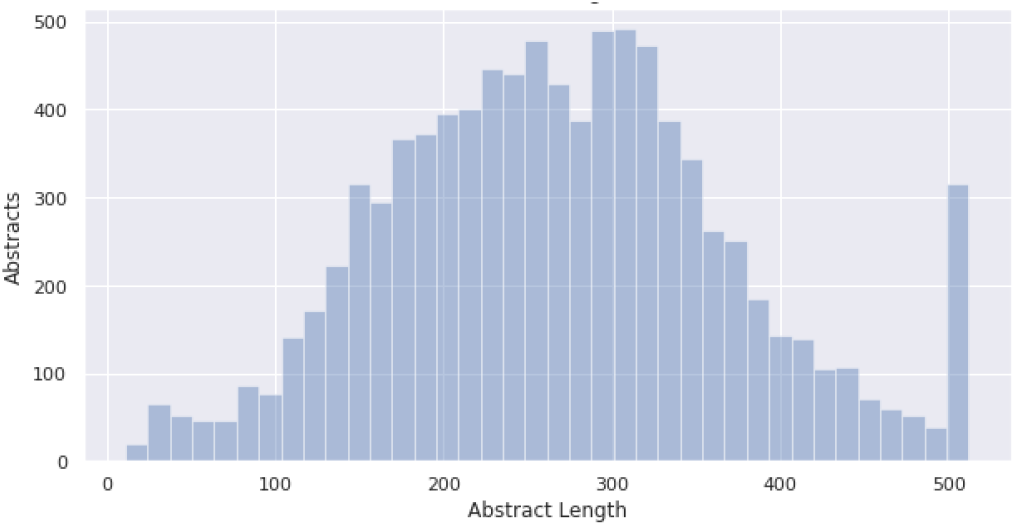
“train and test” Dataset Abstract Character Count Distribution (Post Truncation). Abstracts used for BERT and SciBERT were truncated at a maximum length of 512. Most abstracts were 200-400 characters long in the train/test dataset. Roughly 3OO abstracts needed to be truncated.

**Fig S2.**
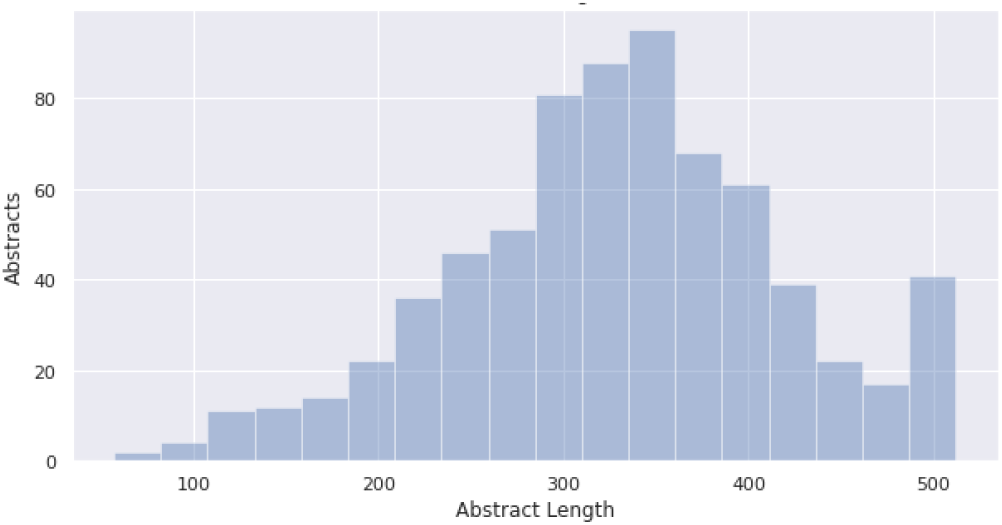
“evaluation” Dataset Abstract Character Count Distribution (Post Truncation). Abstracts used for BERT and SciBERT were truncated at maximum length of 512 characters. Majority of abstracts were 300-400 characters long in the evaluation dataset. Only 40 abstracts needed to be truncated.

**Fig S3.**
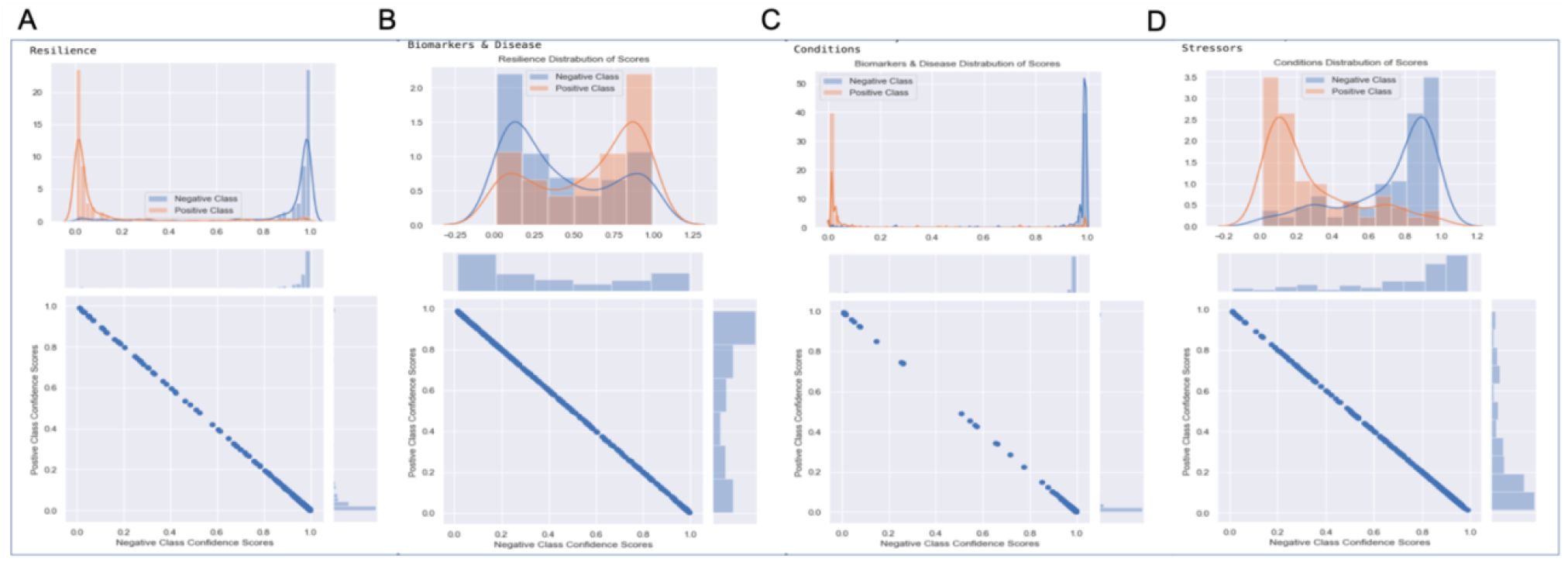
Confidence value distributions of predictions by SciBERT from the “evaluation” dataset. Concepts the model performed well on during the training, such as the Resilience and Conditions categories, had more extreme distributions of scores. Lower scoring categories, such as Biomarkers & Diseases and Stressors, had lower confidence predictions. This suggests that during Artificial Intelligence (AI)/Human augmentation, these categories would need more manual intervention.

**Fig S4.**
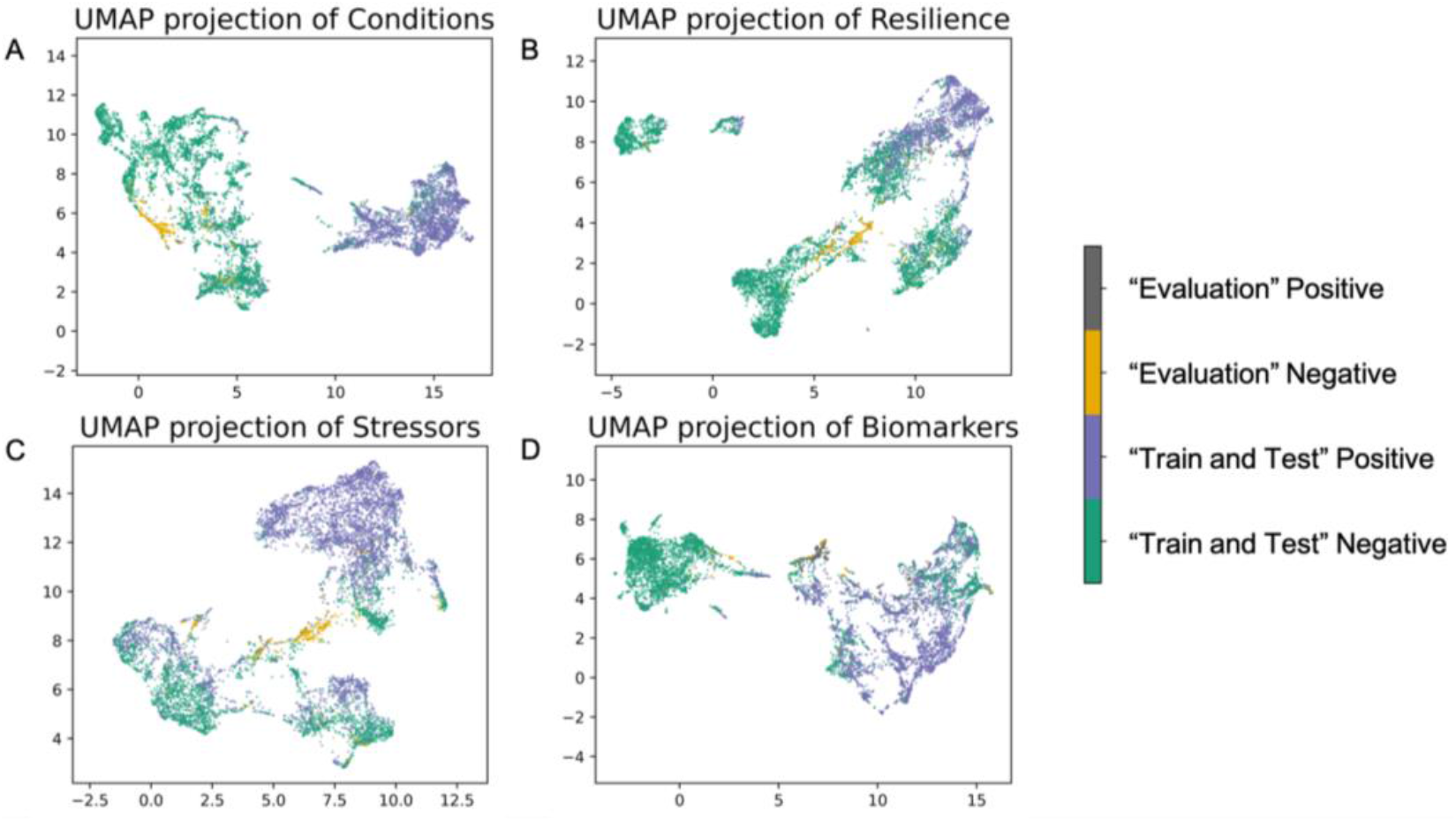
UMAP of embeddings from SciBERT of the “Train and Test” Dataset and subset (*n*=360) of the “Evaluation” Dataset. There was consistent clustering of positive and negative classes. The evaluation data set, overall, cluster with their respective test/train classes suggesting the ability of the model to incorporate novel corpuses of relevant academic literature.

**Table S4.** Training loop times and prediction times for SciBERT. Average prediction time was calculated by measuring how long it took to predict the entire evaluation set and dividing by the length of the evaluation set.

| Category | “Train and Test” dataset training time (seconds) | “Evaluation” dataset prediction time (seconds) | Average time for a single prediction (seconds) |
| --- | --- | --- | --- |
| Resilience | 2128.44 | 13.67 | 0.0189 |
| Biomarkers & Disease | 2103.46 | 13.52 | 0.0187 |
| Conditions | 2103.83 | 13.52 | 0.0187 |
| Stressors | 2107.16 | 13.51 | 0.0187 |

### Trained Reviewers

The consistency and performance of the trained reviewers on meta-concepts varied on sensitivity, specificity, accuracy, and F_1_ scores (Fig S5). Reviewer performance did not correlate with their education, age, or expertise on the scientific domain. Individual reviewer performance was also not consistent across meta-concepts. Even with extensive training the reviewers’ performance was neither 100% accurate nor consistent across meta-concepts compared to ground truth. ML models can perform on par with or better than trained human reviewers. By combining ML and human reviewer efforts as outlined by our human augmentation protocol, we can improve the precision of the abstract review and inclusion process to achieve close to untrained domain expert accuracy and precision.

It is important note that the reviewers were trained and quality-controlled on the “train and test” (psychiatry) dataset and were not provided any additional guidance for transferring the meta-concepts learned in the “train and test” test dataset to the “evaluation” (oncology) dataset.

**Fig S5.**
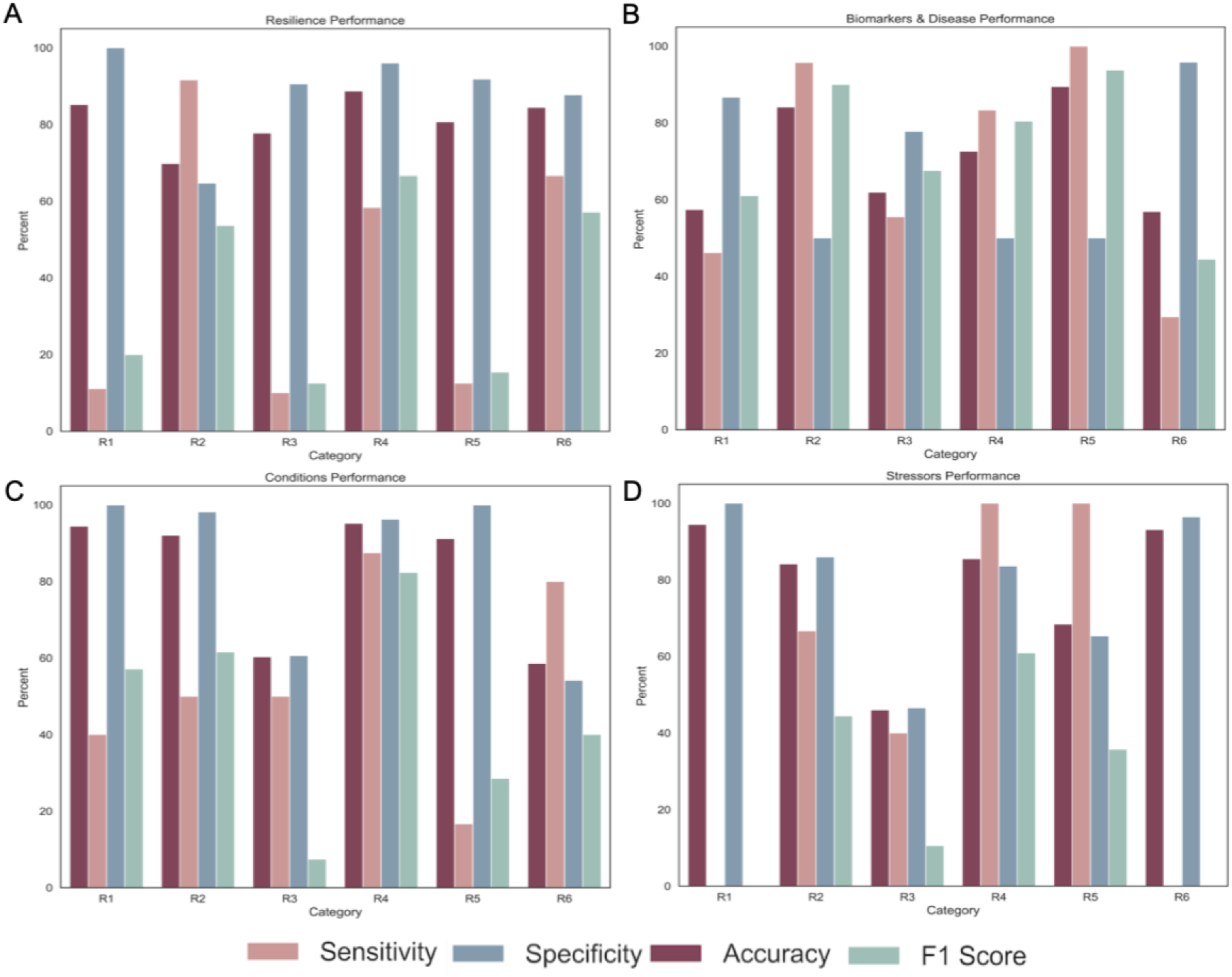
Performance of trained reviewers on the randomly sampled (*n*=36θ) “Evaluation” dataset. The reviewers’ performance were not consistent across education level, age, expertise or concepts.

**Fig S6.**
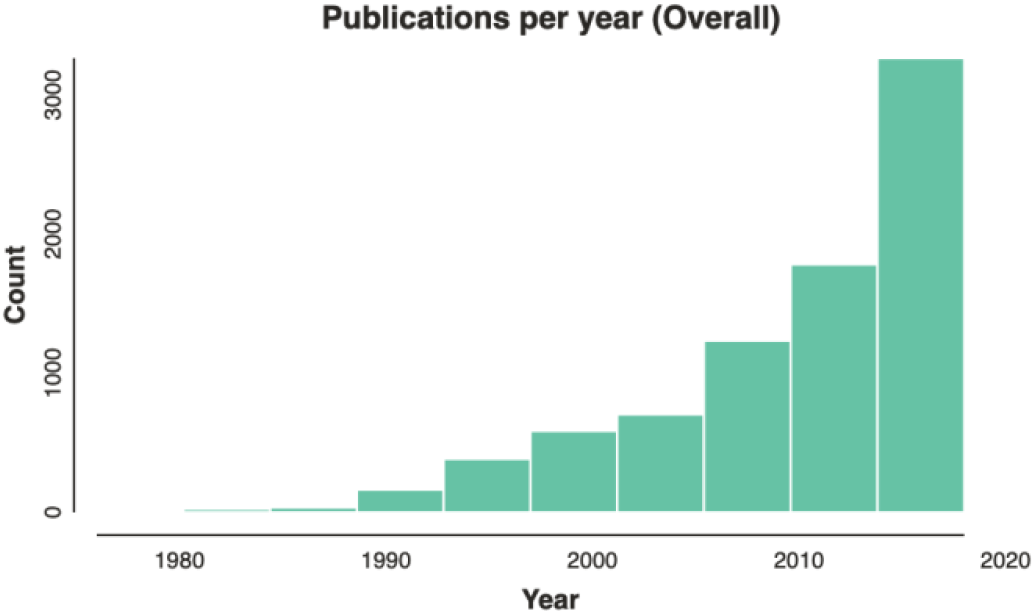
Publications over time in “train and test.” This is based on expert keywords extracted for the meta-analysis examining resilience factors influencing psychopathology development.

**Fig S7.**
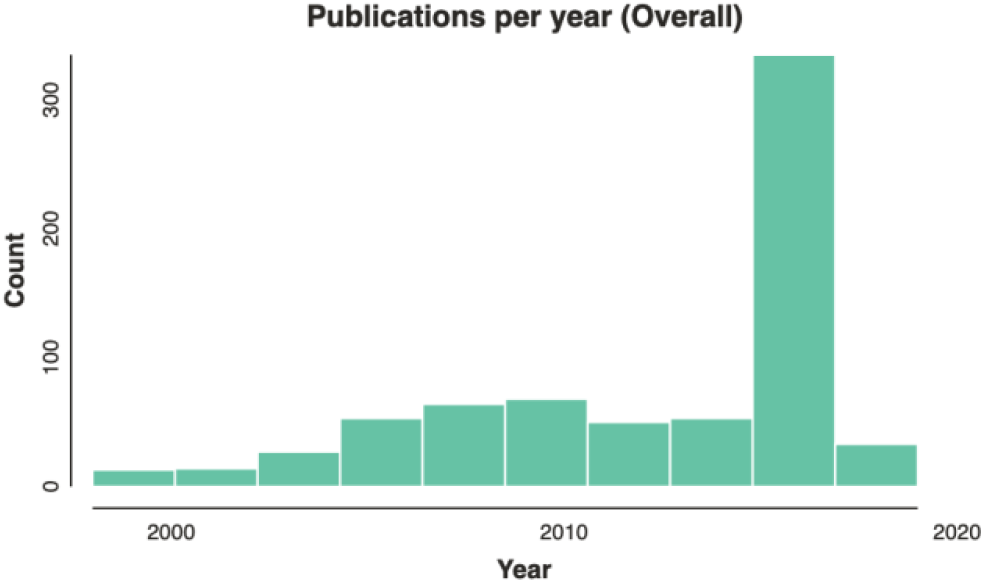
Publications over time in “evaluation” from curated NCI Dataset *n*=720.

**Supplemental Table 3.**
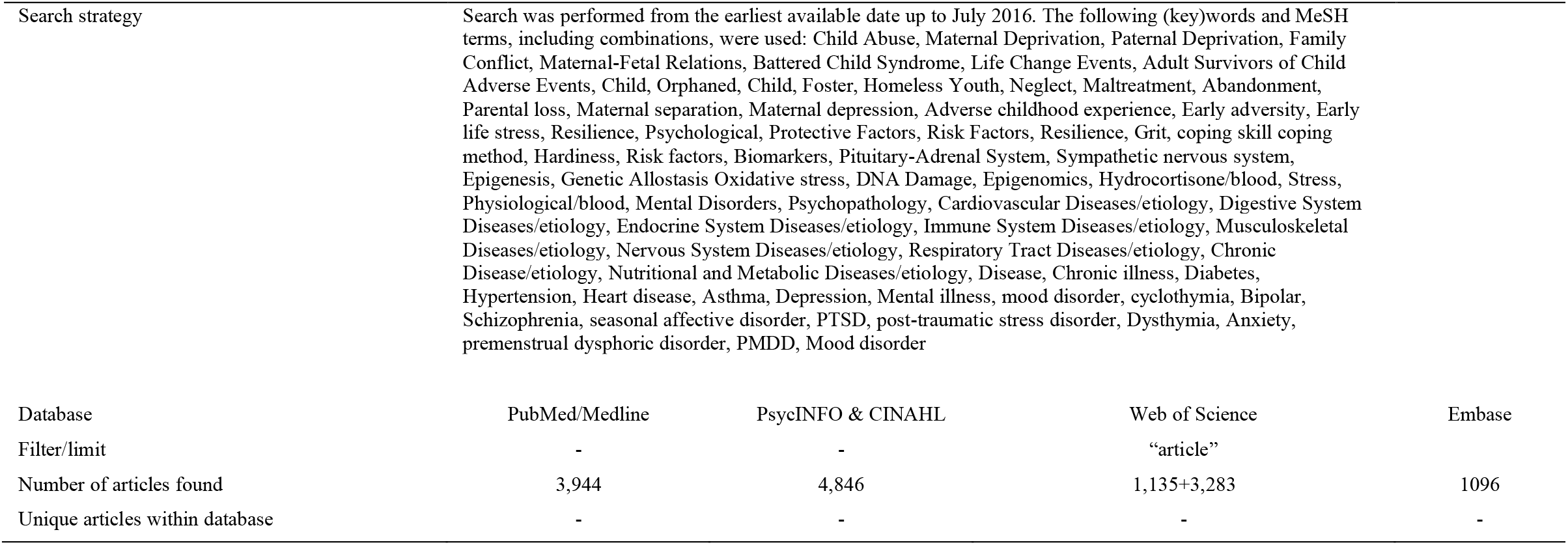
Full search strings

**Supplemental Table 4a.**
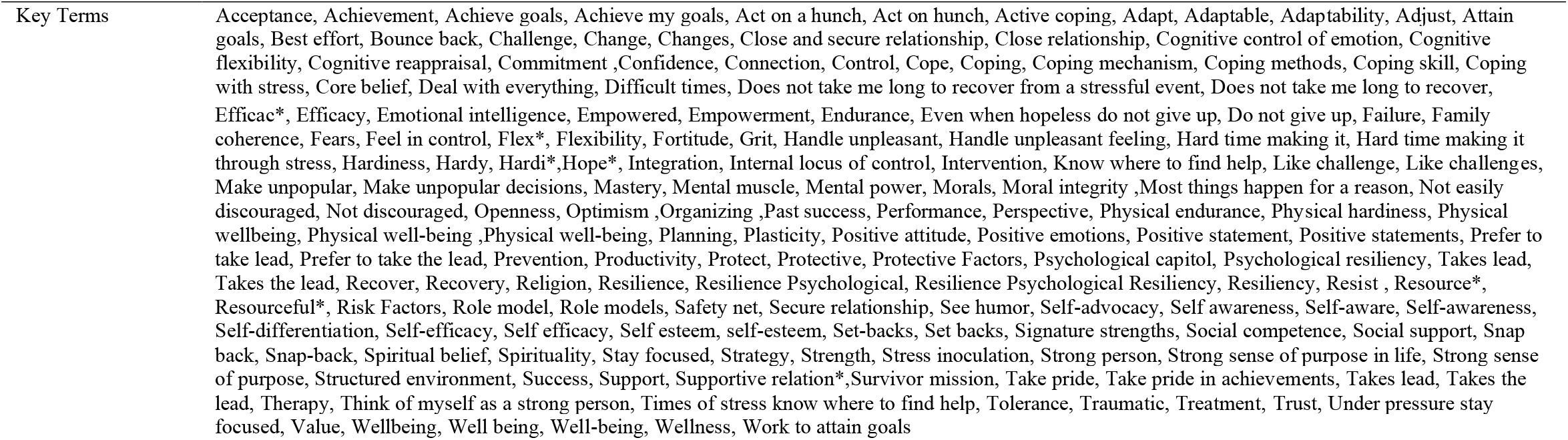
Full key terms for search method: Resilience

**Supplemental Table 4b.**
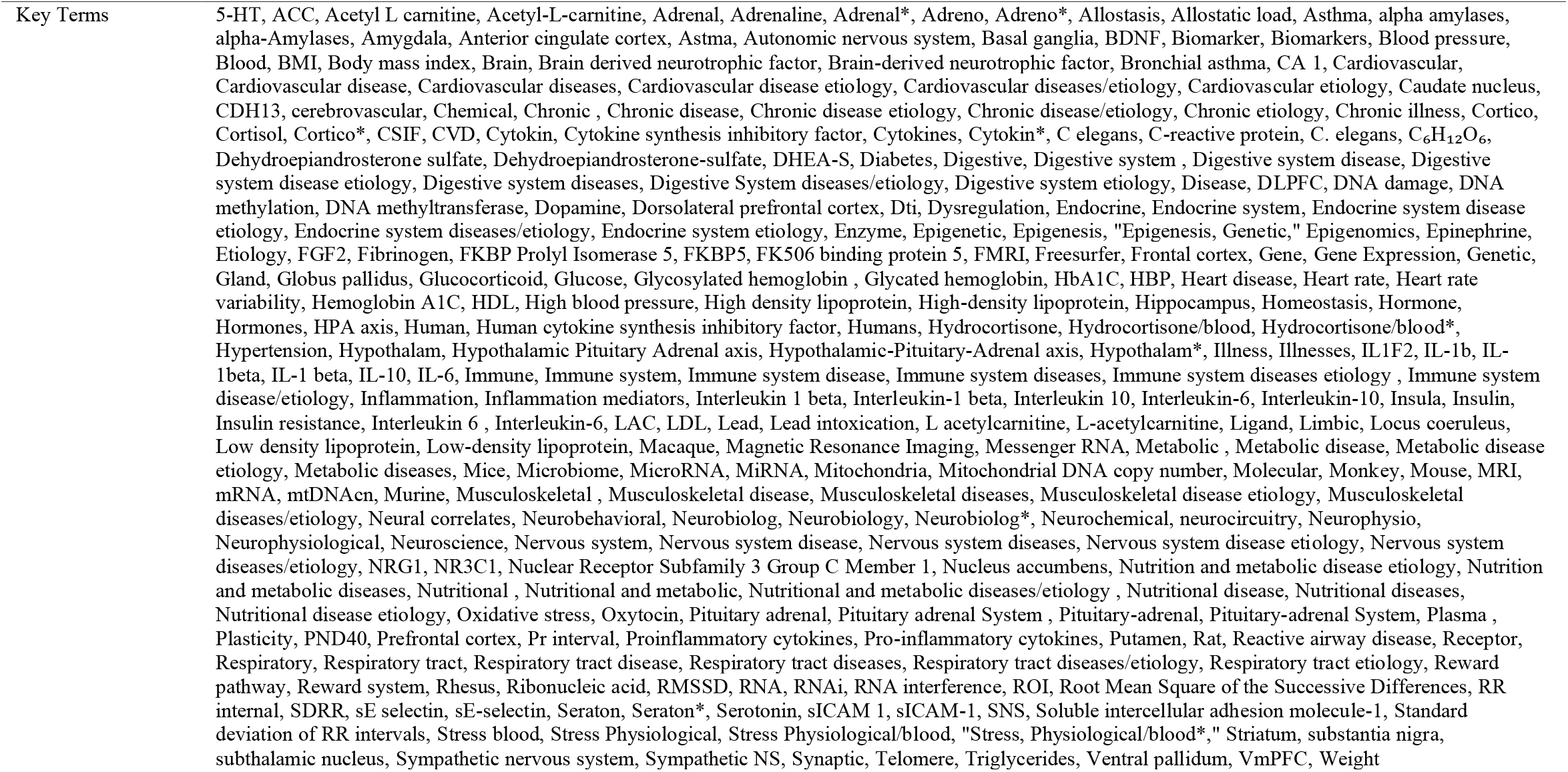
Full key terms for search method: Biomarkers & Disease

**Supplemental Table 4c.**
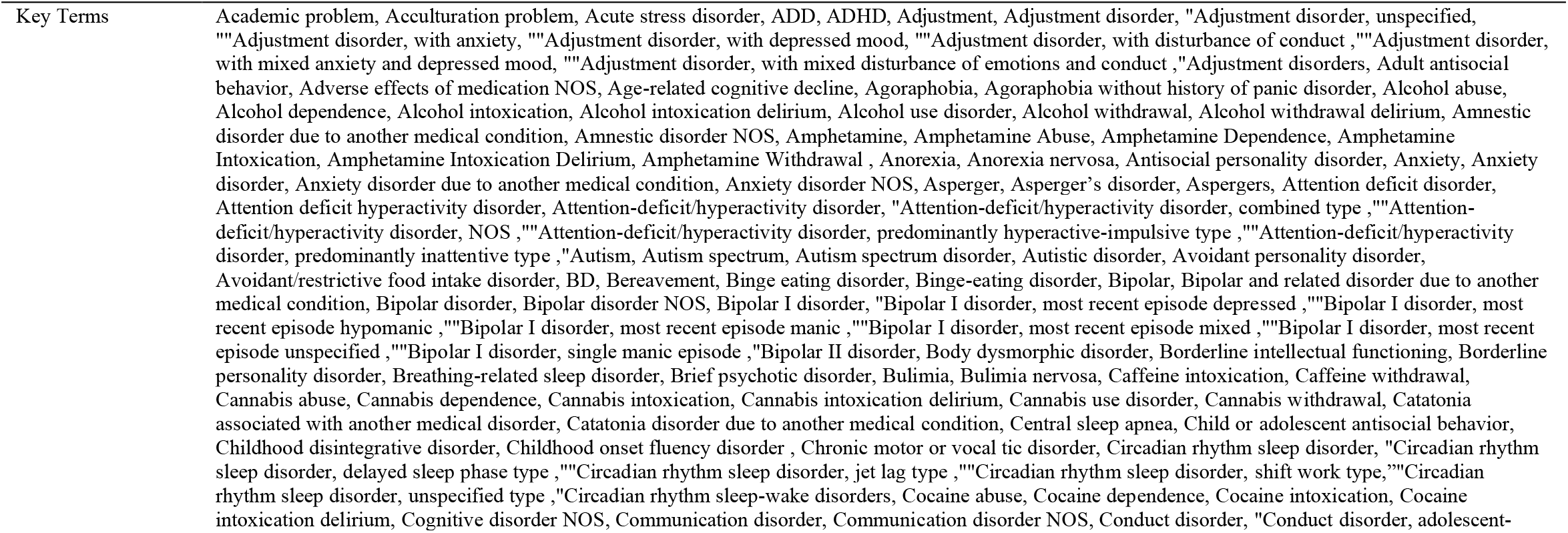
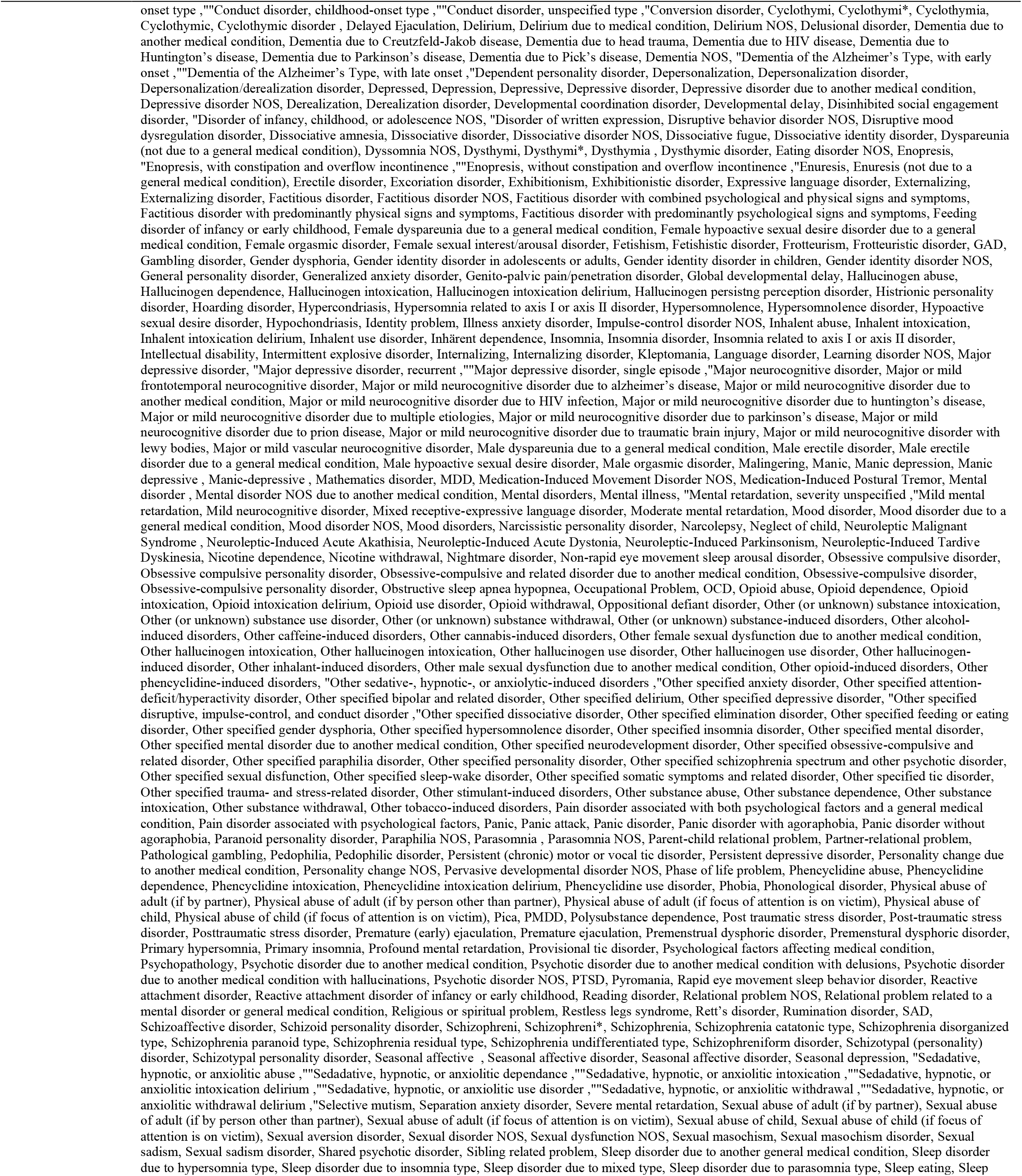
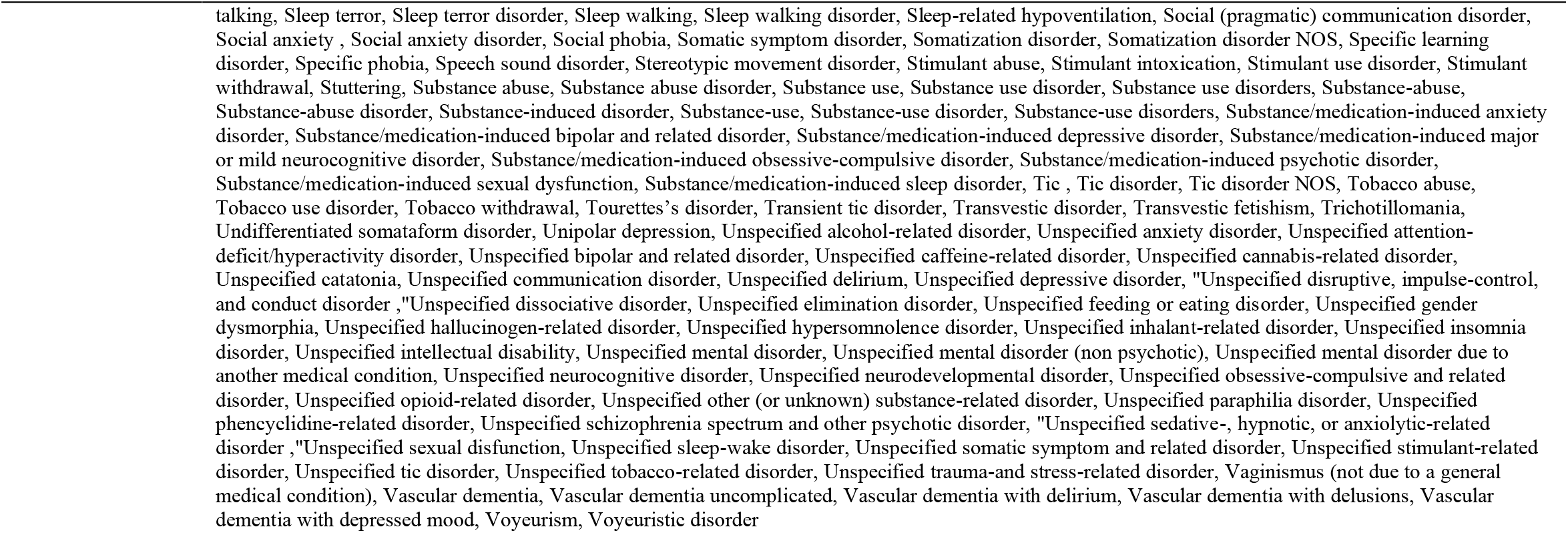
Full key terms for search method: Conditions

**Supplemental Table 4d.**
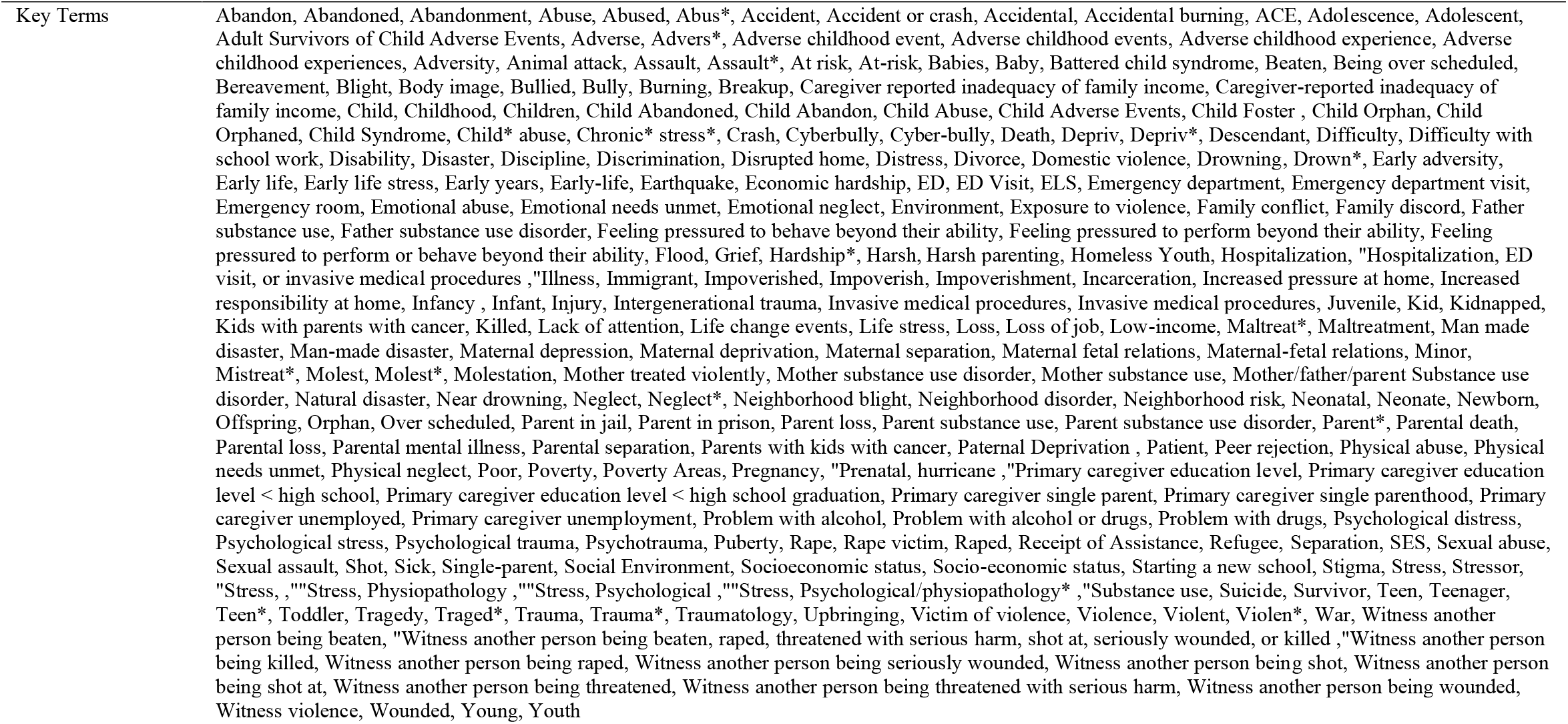
Full key terms for search method: Stressors

